# Phylogeographic analysis of Influenza D virus evolution

**DOI:** 10.1101/2025.09.19.677308

**Authors:** Indrė Blagnytė, Jurgita Markevičiūtė, Gytis Dudas

## Abstract

Influenza D virus (IDV), first identified in 2011 and primarily affecting cattle, has since been detected in a wide range of mammalian hosts and geographic regions. Despite widespread circulation in domestic cattle, the virus’s evolutionary history and global migration dynamics remain poorly understood. This study presents the first comprehensive phylogeographic analysis of IDV using segment 4 (HEF gene) sequences. A time-calibrated Bayesian phylogeographic reconstruction suggests a most recent common ancestor around 1998, with the root most likely in the USA or Japan, although data biases limit definitive conclusions. Generalized linear models (GLMs) were used to assess potential drivers of viral spread, revealing strong associations with cattle trade and shared land borders. However, sequencing efforts heavily influence inferred migration patterns, emphasizing the role of sampling bias in such reconstructions. These findings highlight the mobility of IDV and the critical need for expanded genomic surveillance in underrepresented regions to better detect similar pathogens and understand their transmission dynamics.

## Introduction

Influenza viruses are a group of related viruses in the family *Orthomyxoviridae* and its well-described members include Influenza A virus (IAV), Influenza B virus (IBV), Influenza C virus (ICV), and Influenza D virus (IDV) [fam, 2012, Hause et al., 2014]. *Orthomyxoviridae* genomes consist of segmented linear negative sense single-stranded RNA [fam, 2012]. Even within the influenza virus group segment numbers vary: IAV and IBV have eight segments, ICV and IDV – 7 segments. Proteins encoded by these segments that are common for all influenza viruses include three RNA-dependent RNA polymerase subunits (PB1, PB2 and P3 in ICV and IDV, encoded by segments 1-3), a hemagglutinin (hemagglutinin-esterase fusion-HEF – in ICV and IDV, encoded by segment 4), which is a glycoprotein involved in virus attachment and entry, a nucleoprotein (NP, encoded by segment 5), matrix proteins (M1 and DM2 in IDV, encoded by segment 6), and nonstructural proteins NS1 and NS2 (encoded by segment 7 in ICV and IDV) [Skelton and Huber, 2022].

Influenza A viruses infect humans, causing respiratory disease, while some strains affect other mammals and birds [fam, 2012]. IAV was first isolated from swine in 1930 and from humans in 1933 [Shope, 1931, Smith et al., 1933]. IAV has caused multiple pandemics with death tolls in the millions. The deadliest of these being the 1918 H1N1 pandemic also known as the “Spanish flu”, with estimated deaths around 40 million [Oxford, 2000, Hsieh et al., 2006]. The first cases where recorded in the USA, however the origin country has been disputed. The virus disproportionately affected healthy young adults aged 20–40 [Hsieh et al., 2006]. Many patients died within 24–48 hours due to acute pulmonary edema. The 1957 influenza pandemic, known as the “Asian flu”, caused by subtype H2N2, originated in Hunan, China through genetic reassortment between human and avian influenza strains [Oxford, 2000, Hsieh et al., 2006]. The virus quickly spread worldwide with some death estimates as high as 4 million. It primarily affected infants and the elderly, with pneumonia being the leading cause of death. The 1968 pandemic named “Hong Kong flu” caused by subtype H3N2 resulted in the deaths of an estimated 2 million people. It first appeared in China, spread rapidly through Hong Kong and globally, and caused significant mortality during the 1969–70 winter. While similar in clinical presentation to the 1957 strain, overall mortality was lower, though elderly and young populations remained the most vulnerable. Influenza B first discovered in 1940 mainly infects humans, causing periodic epidemics [Francis Jr, 1940, fam, 2012]. Retrospective analyses of seal sera indicate that the virus likely jumped from humans to seals around the mid-1990s [Osterhaus et al., 2000]. Influenza C was first isolated in 1947 [Taylor, 1949]. It leads to smaller outbreaks in humans and has also been found to infect pigs, dogs, and cattle [fam, 2012, Yuanji et al., 1983, Manuguerra and Hannoun, 1992, Zhang et al., 2018].

While influenza viruses are primarily associated with birds and mammals, newer metagenomic studies have discovered many flu and flu-like viruses in amphibians, fish and jawless vertebrates, suggesting that these aquatic animals might have been among the first hosts of influenza viruses [Parry et al., 2020, Grimwood et al., 2024, Shi et al., 2018, Harding et al., 2022, Petrone et al., 2023, 2024]. Parry et al. [2020] identified three sister lineages to influenza B virus in fish and salamanders and two sister lineages to influenza D virus in amphibians that retained segment conservation and splicing consistent with transcriptional regulation in influenza B and influenza D viruses. Grimwood et al. [2024] discovered respiratory disease-causing influenza and influenza-like viruses in blue antimora and eelpouts (both are fishes). Harding et al. [2022] identified an influenza virus in newts. Shi et al. [2018] documented new influenza viruses in jawless fish, amphibians, and ray-finned fish, with the latter forming a sister group to human influenza B virus, and concluded that host-switching was common during the evolutionary history of influenza viruses. Petrone et al. [2023] identified four divergent, fish-associated influenza-like viruses and suggested that the evolution of orthomyxo- and amnoonviruses was likely shaped by multiple aquatic–terrestrial transitions and substantial host jumps. Influenza-like viruses have also been identified in tunicates, suggesting that influenza viruses emerged prior to the evolution of vertebrates [Petrone et al., 2024].

Influenza D virus was first discovered in pigs with flu-like symptoms in 2011 [Hause et al., 2013]. It was initially classified as a new subtype of influenza C virus because it shared an overall amino acid identity of around 50% with human ICV. Later, however, IDV was classified as a distinct new genus [Hause et al., 2014]. It was also discovered that the primary host for this virus is cattle, rather than swine. The virus causes mild respiratory disease but is also often found in co-infections as part of bovine respiratory disease (BRD) complex caused by a number of unrelated viruses [Ferguson et al., 2016, Ng et al., 2015]. IDV has also been identified in goats, buffalo, giraffes, and wildebeest [Zhai et al., 2017, Molini et al., 2022]. Serological evidence of infection has been found in camelids, goats, sheep, horses, deer, wild boar and cats [Salem et al., 2017, Quast et al., 2015, Nedland et al., 2018, Guan et al., 2022, Gorin et al., 2019, Shen et al., 2025]. Successful experimental infections have been carried out on ferrets, guinea pigs, and mice [Hause et al., 2013, Sreenivasan et al., 2015, Oliva et al., 2020].

A study assessing the prevalence of antibodies against influenza D virus in human sera samples collected in Italy from 2005 to 2017 found seroprevalence ranging from 5.1% to 46.0% depending on the year [Trombetta et al., 2019]. A study by White et al. [2016] found an even higher seroprevalence among people exposed to cattle. A study of 3000 Scottish respiratory samples found no IDV genomes [Smith et al., 2016]. Meanwhile, a study of Malaysian animal workers found the IDV genome in one of 78 samples [Borkenhagen et al., 2018]. With limited research, the effect of IDV on humans remains uncertain although its detection in many artiodactyls suggests broad tropism.

Some phylogenetic analyses of IDV have been conducted, but they have primarily focused on IDV lineages and reassortment. The first two lineages, D/OK and D/660, were identified by Collin et al. in 2015, already noting the frequent reassortment between them. Mekata et al. [2018] found that a Japanese lineage was distinct from the previous two. He et al. [2021] found that IDV has diversified into at least four major lineages (D/OK, D/660, D/Japan, and D/Intermediate), exhibits frequent reassortment, especially involving the D/OK and D/660 strains, and found some geographic clustering. They also estimated the root date to be around 1997. Murakami et al. [2020] identified a second Japanese lineage and renamed them D/Yama2016 and D/Yama2019. In 2021, Huang et al. identified the fifth lineage D/CA2019. Gaudino et al. [2022] found that two primary clades, D/OK and D/660, were circulating in Europe, with D/OK detected earlier and D/660 appearing more recently, coinciding with increased viral diversity and reassortment events. Their root date estimation was around 1995. Limaye et al. [2024] conducted an analysis of all known IDV genomes. They suggested that the earliest ancestors of IDV likely emerged in 1997-1998, with the D/OK lineage emerging in 2005. The study also confirmed a significantly higher substitution rate in IDV than in ICV and identified multiple sub-populations within the D/OK lineage. There have been very few studies examining historical samples for IDV, to corroborate the calculated root dates, however a serological study has shown that IDV has circulated in cattle since at least 2003 [Luo et al., 2017].

Given the lack of knowledge regarding IDV’s origins, both geographically and temporally, as well as limited understanding of how it migrates between countries, we perform a comprehensive Bayesian analysis of publicly available IDV HEF sequences. We carry out Bayesian phylogeographic analyses incorporating various administrative, economic, geographic, demographic and sequencing efforts as correlates of viral migration whilst integrating over phylogenetic uncertainty.

## Methodology

### Sequence data and preprocessing

All influenza D sequences were taken from NCBI Genbank. Only the sequences of segment 4 coding for the HEF gene were chosen, representing the most sequenced segment of the virus. Sequences mislabeled and unlabeled in terms of segment were assigned the correct segment according to the sequence description. Only the sequences that were at least 80% of the expected CDS length were chosen. Any sequences with a percentage of ambiguous nucleotides exceeding 1% were eliminated. Sequences were aligned using G-INS-i method with MAFFT version 7 [Katoh and Standley, 2013] and cropped to only retain the CDS.

### Predictor data

Demographic and economic data was obtained from FAOSTAT and weather data was obtained from Climate Change Knowledge Portal (CCKP) (Table 1). The distances between capitals were great circle distances calculated from geographic coordinates by K. S. Gleditsch [Gleditsch, 2025]. The data for each predictor was standardized (such that the mean of the matrix is zero and variance is one) [Lemey et al., 2014] and log-transformed adding a small offset to datasets with zero values. For each predictor a matrix was constructed with origin country in the rows and destination countries in the columns (Figure S4). For non-symmetrical matrices, two transposed versions were used, each associated with either origin or destination countries. Dummy variables for shared continent and land borders were created manually.

**Table 1.** Data sources used in the study to derive country-level GLM predictor matrices.

| Dataset | Source |
| --- | --- |
| Land Use – Land under agricultural use (%) | FAOSTAT [FAO, 2025a] |
| Trade – Live cattle exports (detailed trade matrix) | FAOSTAT [FAO, 2025b] |
| Trade – Live swine exports (detailed trade matrix) | FAOSTAT [FAO, 2025g] |
| Livestock – Cattle stock | FAOSTAT [FAO, 2025c] |
| Livestock – Swine stock | FAOSTAT [FAO, 2025h] |
| Population – Total population | FAOSTAT [FAO, 2025e] |
| Population – Rural population | FAOSTAT [FAO, 2025f] |
| Macro Indicators – GDP per capita (Value US\$ per capita) | FAOSTAT [FAO, 2025d] |
| Monthly Climatology of Average Mean Surface Air Temperature (1991–2023) | CCKP [World Bank, 2023b] |
| Monthly Climatology of Precipitation (1991–2023) | CCKP [World Bank, 2023a] |
| Distances Between Capital Cities (CAPDIST) dataset | K. S. Gleditsch [Gleditsch, 2025] |

### Exploratory data analysis

An initial phylogenetic tree was inferred using IQ-Tree 2 [Minh et al., 2020]. To find outliers, a root-to-tip regression was performed with TempEst 1.5.3 [Rambaut et al., 2016] and sequences too far from the regression line were eliminated (Figure S1).

Predictor colinearity was examined and predictors were iteratively removed according to pairwise correlation (Figure S2) and suspected importance until none showed a variance inflation factor (VIF) exceeding 5 (Figure S3). The predictors removed from the final analysis were rural population percentage (strongly inversely correlated with GDP per capita) and total human population (strongly correlated with cattle population).

### Bayesian and GLM analysis

BEAST v1.10.4 [Suchard et al., 2018] was used for the Bayesian analysis. Three independent MCMC runs were performed, each with a chain length of 100 million, logging parameters every 10,000 states.

For nucleotide evolution, the HKY substitution model [Hasegawa et al., 1985] was applied with estimated base frequencies and gamma-distributed rate variation among sites [Yang, 1996] (four discrete categories). The analysis included two partitions based on codon positions 1+2 and 3 (i.e. the SRD06 site model [Shapiro et al., 2006]). Additional model parameters were: an uncorrelated relaxed clock with log-normally distributed rates [Drummond et al., 2006], and a Bayesian SkyGrid tree prior with 50 time points and a cutoff of 25 years [Gill et al., 2013, Drummond et al., 2002]. A GLM model was set up [Lemey et al., 2014], incorporating previously described predictor matrices. Bayes factors were calculated using the formula:

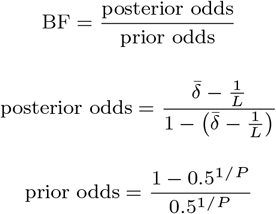

where:

- BF is the Bayes Factors
- 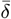 the proportion of MCMC samples in which the predictor is included
- *L* is the total number of MCMC samples (MCMC length), where 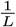 is a small correction factor to avoid division by 0 when computing posterior odds for predictors that are always included in the model, *i.e*. 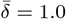
- *P* is the number of predictors in the model

The burn-in size (10 million which represents 10% of each run) for the MCMC results was determined using Tracer v1.7.2 [Rambaut et al., 2018]. The three runs were then combined using LogCombiner v1.10.4, and a Maximum Clade Credibility (MCC) tree was generated using common ancestor heights with TreeAnnotator v1.10.4.

## Results

A detailed phylodynamic analysis of the currently available influenza D data and the factors influencing its evolution has not been previously conducted. To address this gap, a comprehensive phylogeographic reconstruction was carried out here to characterise the evolutionary history and migration patterns of IDV.

In the maximum clade credibility (MCC) tree, all 5 known lineages of IDV were visible (Fig. 1). The root was inferred to be in the USA, with a posterior probability of 0.62, with the second most likely country of origin being Japan (0.31) and remaining sampled countries having negligible probabilities. The mean posterior root date was calculated as 1997.86 (95% HPD: 1991.37-2003.18), with the median being 1998.42, suggesting that all currently sequenced IDV HEF sequences are descended from an ancestor in the USA around the year 1998. However, the probability density function is rather spread out, with just the inter-quartile range spanning around 3.5 years.

**Fig. 1:**
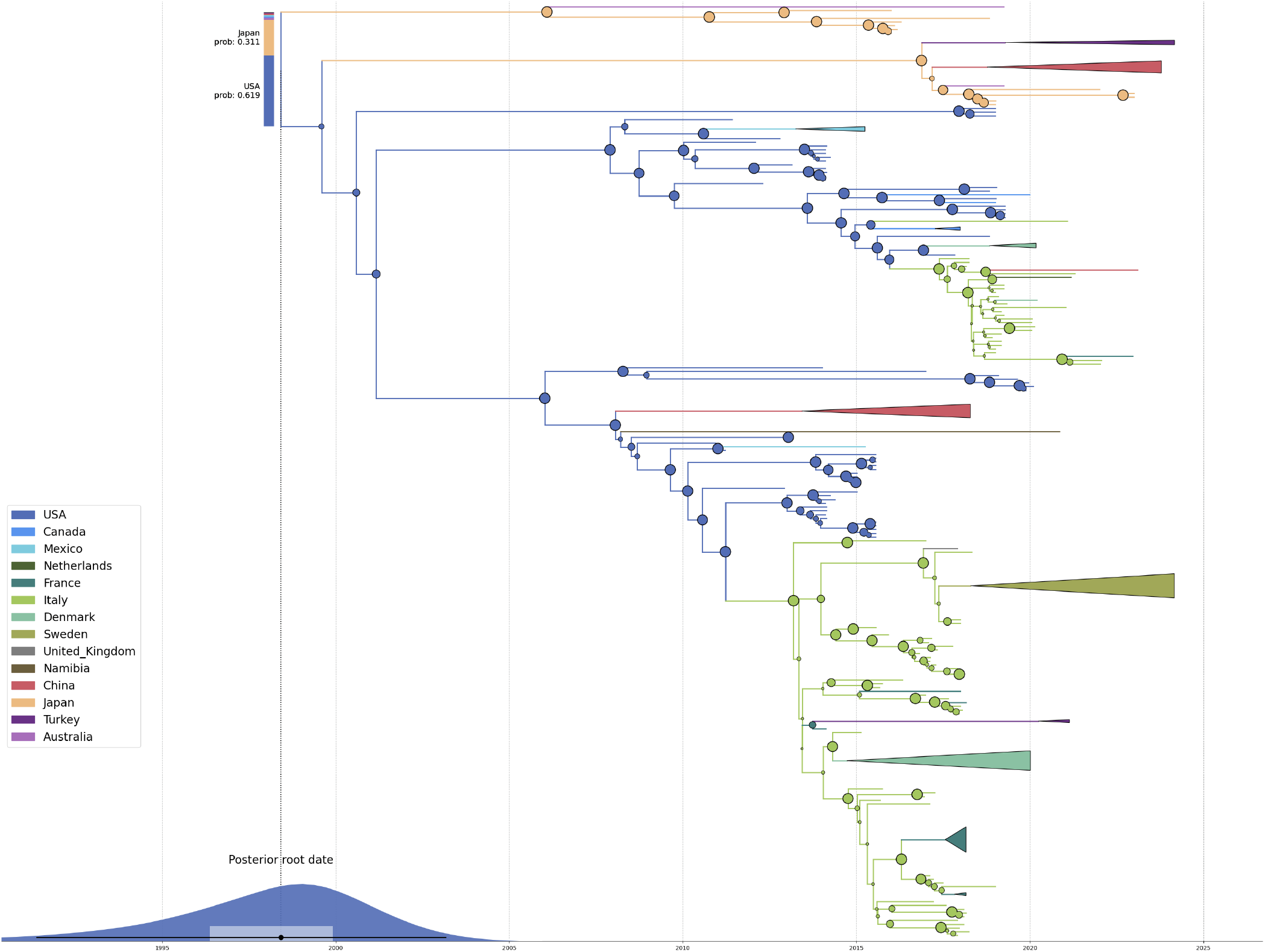
MCC tree of influenza D virus HEF segments. Branches are coloured by inferred country (legend on the left) with some clades of closely related sequences from the same country collapsed. Circles at nodes are proportional to their posterior probabilities. The inferred date of the root is marked with a blue kernel density estimate at the bottom where the mean estimate is marked with a black dot. The bar next to the root indicates the posterior probabilities of each inferred ancestral state at the root.

The initial state-space GLM showed two significant predictors-land borders and cattle trade for the destination country (table 2). The trade matrices were created using export data, meaning that the destination matrix shows the cattle imports into the origin country. Both predictor Bayes factors exceeded 10, showing very strong evidence for inclusion into the GLM model, with the cattle trade predictor showing stronger evidence with a Bayes factor of 38.8. Both predictors showed a positive linear relationship with virus migrations, with land borders showing a higher regression weight (2.22) than trade (0.65). Unlike cattle trade, swine trade did not show any evidence for inclusion into the model, demonstrating that while cattle trade appears to have an impact on virus migration, swine trade does not.

**Table 2.** GLM results – model predictors with mean conditional beta coefficients (effect size) and their 95% highest probability density intervals, indicator means (inclusion probability), and Bayes factors (statistical support). Bold rows highlight predictors with strong statistical support (BF*≥*3).

| No. | Predictor | Mean Conditional Beta Coefficient | 95% HPD Interval of Beta Coeff. | Indicator Mean | Bayes Factor |
| --- | --- | --- | --- | --- | --- |
| 1 | agro_land_perc_destination_logtransform | 0.16 | [-0.14, 0.50] | 0.00 | 0.01 |
| 2 | agro_land_perc_origin_logtransform | 0.57 | [-1.56, 3.39] | 0.00 | 0.09 |
| 3 | capital_distance_km_offsetlog | -1.38 | [-2.33, -0.49] | 0.05 | 1.45 |
| **4\*** | **continent** | **1.41** | **[0.65, 2.23]** | **0.22** | **7.74** |
| 5 | GDP_per_capita_destination_logtransform | 0.06 | [-0.22, 0.57] | 0.00 | 0.01 |
| 6 | GDP_per_capita_origin_logtransform | -0.13 | [-1.35, 1.38] | 0.00 | 0.07 |
| **7\*** | **land_borders** | **1.40** | **[0.53, 2.18]** | **0.14** | **4.33** |
| 8 | total_cattle_stock_destination_logtransform | 0.08 | [-0.26, 0.47] | 0.00 | 0.01 |
| 9 | total_cattle_stock_origin_logtransform | 0.05 | [-0.41, 0.42] | 0.00 | 0.02 |
| 10 | total_swine_stock_destination_logtransform | 0.18 | [-0.07, 0.51] | 0.00 | 0.03 |
| 11 | total_swine_stock_origin_logtransform | 0.19 | [-0.91, 1.32] | 0.00 | 0.06 |
| 12 | swine_trade_exp_destination_offsetlog | 0.29 | [0.08, 0.51] | 0.01 | 0.17 |
| 13 | swine_trade_exp_origin_offsetlog | 0.41 | [0.05, 0.78] | 0.00 | 0.11 |
| 14 | yearly_precipitation_destination_logtransform | 0.06 | [-0.26, 0.40] | 0.00 | 0.02 |
| 15 | yearly_precipitation_origin_logtransform | 0.78 | [-2.08, 2.49] | 0.01 | 0.22 |
| 16 | mean_temperatures_destination_logtransform | 0.07 | [-0.37, 0.31] | 0.00 | 0.01 |
| 17 | mean_temperatures_origin_logtransform | 0.72 | [-1.23, 2.95] | 0.00 | 0.07 |
| **18\*** | **num_of_seq_origin_logtransform** | **3.15** | **[1.17, 5.33]** | **1.00** | **11.60** |
| 19 | num_of_seq_destination_logtransform | 0.51 | [-0.03, 0.94] | 0.01 | 0.26 |

**Table 3.** GLM results for exploring the significance of the number of sequences. Includes mean conditional beta coefficients and their 95% highest probability density intervals, indicator means, and Bayes factors. Bold rows highlight predictors with statistical support (BF*≥*3). In this analysis cattle trade predictors were included as predictors (underlined) and swine trade predictors were excluded.

| No. | Predictor | Mean Conditional Beta Coefficient | 95% HPD Interval of Beta Coeff. | Indicator Mean | Bayes Factor |
| --- | --- | --- | --- | --- | --- |
| 1 | agro_land_perc_destination_logtransform | 0.07 | [-0.25, 0.41] | 0.00 | 0.03 |
| 2 | agro_land_perc_origin_logtransform | 0.34 | [-1.14, 2.23] | 0.00 | 0.06 |
| 3 | capital_distance_km_offsetlog | -1.37 | [-2.30, -0.37] | 0.05 | 1.31 |
| **4\*** | **continent** | **1.40** | **[0.64, 2.25]** | **0.20** | **6.79** |
| 5 | GDP_per_capita_destination_logtransform | 0.02 | [-0.38, 0.41] | 0.00 | 0.02 |
| 6 | GDP_per_capita_origin_logtransform | -0.08 | [-1.38, 1.25] | 0.00 | 0.08 |
| **7\*** | **land_borders** | **1.40** | **[0.55, 2.18]** | **0.13** | **3.88** |
| 8 | total_cattle_stock_destination_logtransform | 0.15 | [-0.07, 0.62] | 0.00 | 0.01 |
| 9 | total_cattle_stock_origin_logtransform | 0.10 | [-0.32, 0.54] | 0.00 | 0.02 |
| 10 | total_swine_stock_destination_logtransform | 0.10 | [-0.51, 0.57] | 0.00 | 0.01 |
| 11 | total_swine_stock_origin_logtransform | 0.25 | [-0.96, 1.57] | 0.00 | 0.06 |
| 12 | <u>cattle_trade_exp_destination_offsetlog</u> | 0.43 | [0.19, 0.70] | 0.08 | 2.49 |
| 13 | <u>cattle_trade_exp_origin_offsetlog</u> | 0.41 | [0.08, 0.80] | 0.01 | 0.27 |
| 14 | yearly_precipitation_destination_logtransform | 0.07 | [-0.40, 0.52] | 0.00 | 0.01 |
| 15 | yearly_precipitation_origin_logtransform | 1.01 | [-1.69, 2.46] | 0.01 | 0.27 |
| 16 | mean_temperatures_destination_logtransform | 0.00 | [-0.42, 0.85] | 0.00 | 0.01 |
| 17 | mean_temperatures_origin_logtransform | 0.47 | [-1.49, 2.96] | 0.00 | 0.09 |
| **18\*** | **num_of_seq_origin_logtransform** | **3.12** | **[1.11, 5.34]** | **0.99** | **4351.19** |
| 19 | num_of_seq_destination_logtransform | 0.56 | [0.18, 1.00] | 0.01 | 0.32 |

Examining individual ancestry plots reveals that many strains jump across three different countries in two continents (Fig. S5C-E), and in the case of Fig. S5B, the HEF sequence from China (D/bovine/CHN/JY3002/2022) migrations across three continents are inferred. This shows how quickly and broadly the virus is able to spread across geographically distant areas. In the case of Fig S5A, the plot also shows a lot of uncertainty, demonstrating the lack of data from many countries that creates the risk of biased results.

Analysing the migration map (Fig. S6), multiple trans-Atlantic transitions can be seen showing the global spreading potential that IDV has. It is also evident that most migrations originate in either the US or Italy. Since these countries are also the ones with the most available IDV sequences, we explored whether these sequencing biases affect our phylogeographic reconstruction.

To explore the effect of the amount of data from each country on virus migration reconstruction, another GLM was set up using a predictor matrix with number of sequences available for each country as origin and destination predictors. Due to multi-collinearity limitations, two GLMs were set up. One excluding pig trade and one excluding cattle trade data. In the new analyses, the one including cattle trade and the one with swine trade, the number of sequences for the origin countries shows by far the largest inclusion evidence with the Bayes factor in the thousands (4351.19 and 9411.60, respectively) and the largest effect size (3.12 and 3.15). The land borders predictor retains evidence for inclusion, though with a smaller effect size (1.40 in both cases). The continent predictor gains moderate inclusion evidence (BF of 6.79 and 7.74) with effect sizes similar to the land border predictor (1.40 and 1.41). Interestingly, the cattle trade predictor Bayes factor falls below the threshold for inclusion (2.49 – weak evidence for inclusion) even though it doesn’t show a large pairwise correlation with the number of sequences predictor. Although the Bayes factor for this predictor is below the conventional threshold for strong evidence, its conditional beta coefficient is consistently positive with a 95% HPD interval excluding zero. This suggests that, when included in the model, the predictor tends to have a reliably positive effect, even if the overall evidence for its inclusion is limited.

Overall, available influenza D virus sequence data indicate relatively fast and broad spread with jumps between multiple continents. The root is inferred to be in the USA, but because of the disproportionate amount of data from the USA, Asian origins cannot be rejected. GLM results indicate that the amount of available sequence data per country has a substantial influence on the inferred viral migration patterns. Other contributing factors include geographical relationships and potentially cattle trade.

## Discussion

This study presents the first comprehensive phylogeographic analysis of Influenza D virus (IDV), integrating evolutionary history with potential drivers of viral migration. The results provide new insights into the global spread of IDV and the factors influencing its transmission dynamics.

The inferred root date of 1998 aligns closely with Gaudino et al. and Limaye et al. analyses. While the inferred location of the root appears to be in the USA, it also has been demonstrated that the analysis is strongly impacted by the amount of data from each country, with the USA having the largest amount of sequence data available. This suggests that the USA may not be the location where currently sampled influenza D virus HEF diversity originated. With the amount of diversity of Influenza D virus that Asia exhibits, and considering the calculated 0.31 probability of the root being in Japan, it is possible that the root of the tree is indeed actually in Asia and not in the USA as the analysis suggests. Broader and representative sequencing efforts may easily identify alternative country of origin that is a different Asian country than Japan, considering that currently only Japan and China have any significant number of sequences available. Historical samples may prove particularly enlightening too.

In some cases, cattle trade data, specifically cattle imports to the origin country, showed evidence for inclusion into the GLM model. This demonstrates that international trade of the primary host species plays a significant role in the cross-border movement of IDV. In contrast, despite the known presence of IDV in pigs, swine trade did not show any meaningful effect on inferred viral migration patterns. However, the analysis also highlights an important limitation: the strong influence of data availability on migration inference. Countries with extensive sequencing efforts, such as the United States and Italy, appear as central hubs in the migration network. This likely reflects sampling bias rather than true epidemiological centrality, as countries with limited or no sequence data are effectively excluded from the reconstruction.

RNA viruses are known for crossing species barriers easily, thus posing a significant threat to human health. One such example of zoonotic potential is the 2001 discovery of human metapneumovirus (hMPV) in children with respiratory illness in the Netherlands [Van den Hoogen et al., 2001]. Initially identified as closely related to the avian pneumovirus, hMPV was shown through retrospective serological studies to have been circulating in humans for at least 50 years. It is associated with both upper and lower respiratory infections and particularly affects infants, the elderly, and those with chronic health conditions [Williams et al., 2004, Edwards et al., 2013, Boivin et al., 2002, 2007, Williams et al., 2005, van den Hoogen et al., 2003]. Similarly, highly pathogenic avian influenza (HPAI) A(H5N1), first identified in 1996 [Xu et al., 1999], continues to evolve and pose risks to both animal and human health. HPAI A(H5N1) clade 2.3.4.4b is currently the predominant clade and infects both poultry and mammals [Huang et al., 2023]. In early 2024, infections with clade 2.3.4.4b influenza A viruses were detected in dairy cows in the US, with over 1,073 dairy herds affected as of June 2025 [Burrough et al., 2024, U.S. Centers for Disease Control and Prevention, 2025]. Most cow infections are genotype B3.13, with most outbreaks in wild birds and poultry being genotype D1.1 [Zhang et al., 2025]. Human infections have also been recorded, with the majority being caused by genotype B3.13. [Garg et al., 2025]. The zoonotic potential of RNA viruses, such as human metapneumovirus and highly pathogenic avian influenza A(H5N1), highlights the broader relevance of interspecies transmission in shaping viral emergence and spread. Similar to these viruses, influenza D virus — while currently confined largely to cattle and pigs — warrants close monitoring given its capacity to cross species barriers and its detection in multiple hosts to date.

A related argument can be made regarding Tilapia lake virus - first identified following widespread mortalities of wild and farmed hybrid tilapia in Israel during the summer of 2014 [Eyngor et al., 2014]. The virus raises alarm due to the species’ global economic importance as a major source of affordable protein and income for fish farmers [FAO, 2017]. Although the virus was only formally identified in 2013, genomic analyses suggest its emergence occurred as early as 2003–2009 [Thawornwattana et al., 2021]. This case exemplifies how pathogens can remain undetected in agricultural contexts for years, until significant losses affecting economically vital livestock species spur a response. The discovery of Tilapia lake virus mirrors that of influenza D virus, stressing the need for proactive and agnostic surveillance in farmed animals to detect and respond to novel viruses before they become widely established and have a chance to pose a threat to human health.

Furthermore, the recent discovery of a substantial and previously unknown diversity of influenza viruses in fish and amphibians, many of which are more closely related to human-pathogenic influenza A, B and C viruses than each other, as well as consistent inference of currently sampled influenza D diversity dating back approximately 25 years raises questions about its origins. For one, it’s still unclear if influenza D viruses have undergone population turnover since their jump into cattle and correspondingly whether the roughly 1997 estimate of the root date here corresponds to the date of the jump from some reservoir species or population processes that happened cattle-side. Similarly, identifying the reservoir host(s) of influenza D virus prior to its appearance in cattle may significantly shift our understanding of zoonotic risks posed by influenza viruses to humans, particularly if these hosts turn out to be more distantly related to cattle than expected, such as fish or other early-diverging vertebrates.

Overall, the findings accentuate the need for more balanced and widespread genomic surveillance of IDV, particularly in underrepresented regions. Increasing the availability of sequence data from a broader and representative set of countries will be crucial for improving the accuracy of future phylodynamic studies and for understanding the true global ecology of influenza D virus.

## Data availability

All data and code are made publicly available on Github: https://github.com/indrebl/flu d project

## Acknowledgments

G.D. is supported by EMBO installation grant EMBO-IG-5305-2023.

## Supplementary materials

**Fig. S1:**
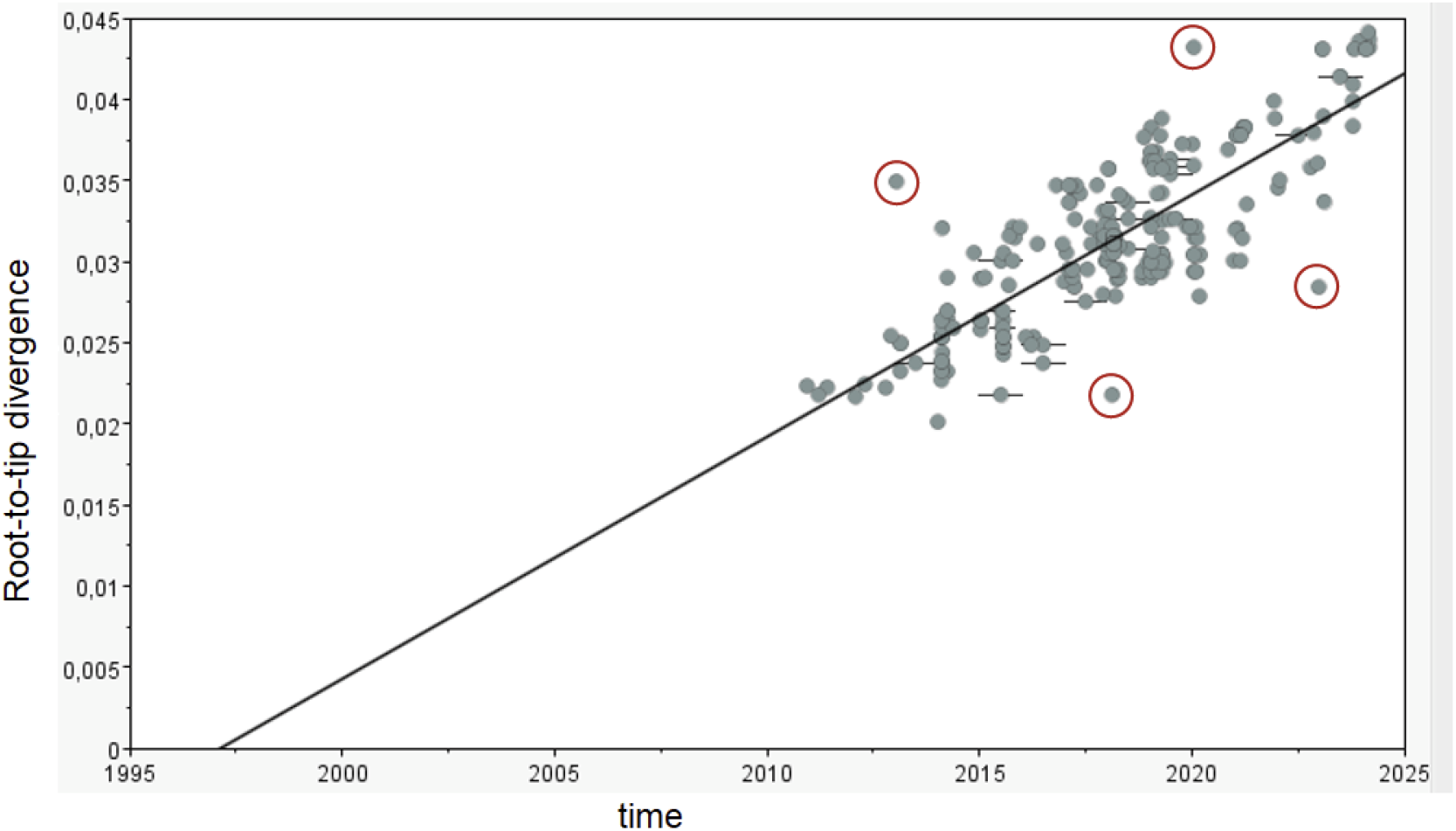
Root-to-tip regression of influenza D HEF maximum likelihood tree. The removed outliers are circled in red.

**Table S1.** GLM results for exploring the significance of the number of sequences. Includes mean conditional beta coefficients and their 95% highest probability density intervals, indicator means, and Bayes factors. Bold rows highlight predictors with strong Bayes factors. In this analysis swine trade predictors were included (underlined) as predictors and cattle trade predictors were excluded.

| No. | Predictor | Mean Conditional Beta Coefficient | 95% HPD Interval of Beta Coeff. | Indicator Mean | Bayes Factor |
| --- | --- | --- | --- | --- | --- |
| 1 | agro_land_perc_destination_logtransform | 0.16 | [-0.14, 0.50] | 0.00 | 0.01 |
| 2 | agro_land_perc_origin_logtransform | 0.57 | [-1.56, 3.39] | 0.00 | 0.09 |
| 3 | capital_distance_km_offsetlog | -1.38 | [-2.33, -0.49] | 0.05 | 1.45 |
| **4\*** | **continent** | **1.41** | **[0.65, 2.23]** | **0.22** | **7.74** |
| 5 | GDP_per_capita_destination_logtransform | 0.06 | [-0.22, 0.57] | 0.00 | 0.01 |
| 6 | GDP_per_capita_origin_logtransform | -0.13 | [-1.35, 1.38] | 0.00 | 0.07 |
| **7\*** | **land_boarders** | **1.40** | **[0.53, 2.18]** | **0.14** | **4.33** |
| 8 | total_cattle_stock_destination_logtransform | 0.08 | [-0.26, 0.47] | 0.00 | 0.01 |
| 9 | total_cattle_stock_origin_logtransform | 0.05 | [-0.41, 0.42] | 0.00 | 0.02 |
| 10 | total_swine_stock_destination_logtransform | 0.18 | [-0.07, 0.51] | 0.00 | 0.03 |
| 11 | total_swine_stock_origin_logtransform | 0.19 | [-0.91, 1.32] | 0.00 | 0.06 |
| 12 | <u>swine_trade_exp_destination_offsetlog</u> | 0.29 | [0.08, 0.51] | 0.01 | 0.17 |
| 13 | <u>swine_trade_exp_origin_offsetlog</u> | 0.41 | [0.05, 0.78] | 0.00 | 0.11 |
| 14 | yearly_precipitation_destination_logtransform | 0.06 | [-0.26, 0.40] | 0.00 | 0.02 |
| 15 | yearly_precipitation_origin_logtransform | 0.78 | [-2.08, 2.49] | 0.01 | 0.22 |
| 16 | mean_temperatures_destination_logtransform | 0.07 | [-0.37, 0.31] | 0.00 | 0.01 |
| 17 | mean_temperatures_origin_logtransform | 0.72 | [-1.23, 2.95] | 0.00 | 0.07 |
| **18\*** | **num_of_seq_origin_logtransform** | **3.15** | **[1.17, 5.33]** | **1.00** | **9411.60** |
| 19 | num_of_seq_destination_logtransform | 0.51 | [-0.03, 0.94] | 0.01 | 0.26 |

**Fig. S2:**
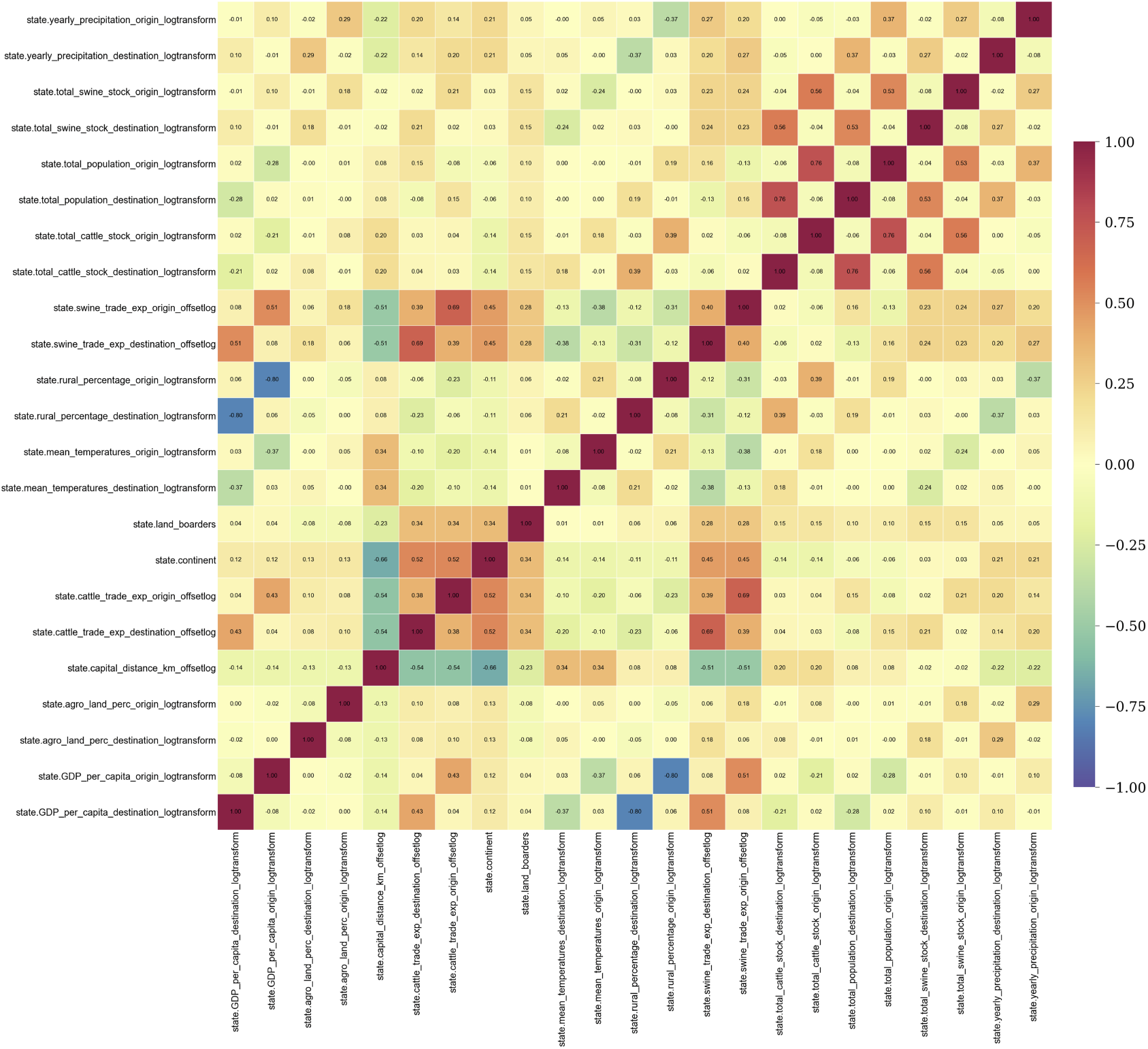
Pairwise Spearman correlations between all predictor variables before variable elimination. Colours indicate magnitude and direction of the correlation (red is strongly positive, blue is strongly negative) with correlation coefficient added as text to each cell.

**Fig. S3:**
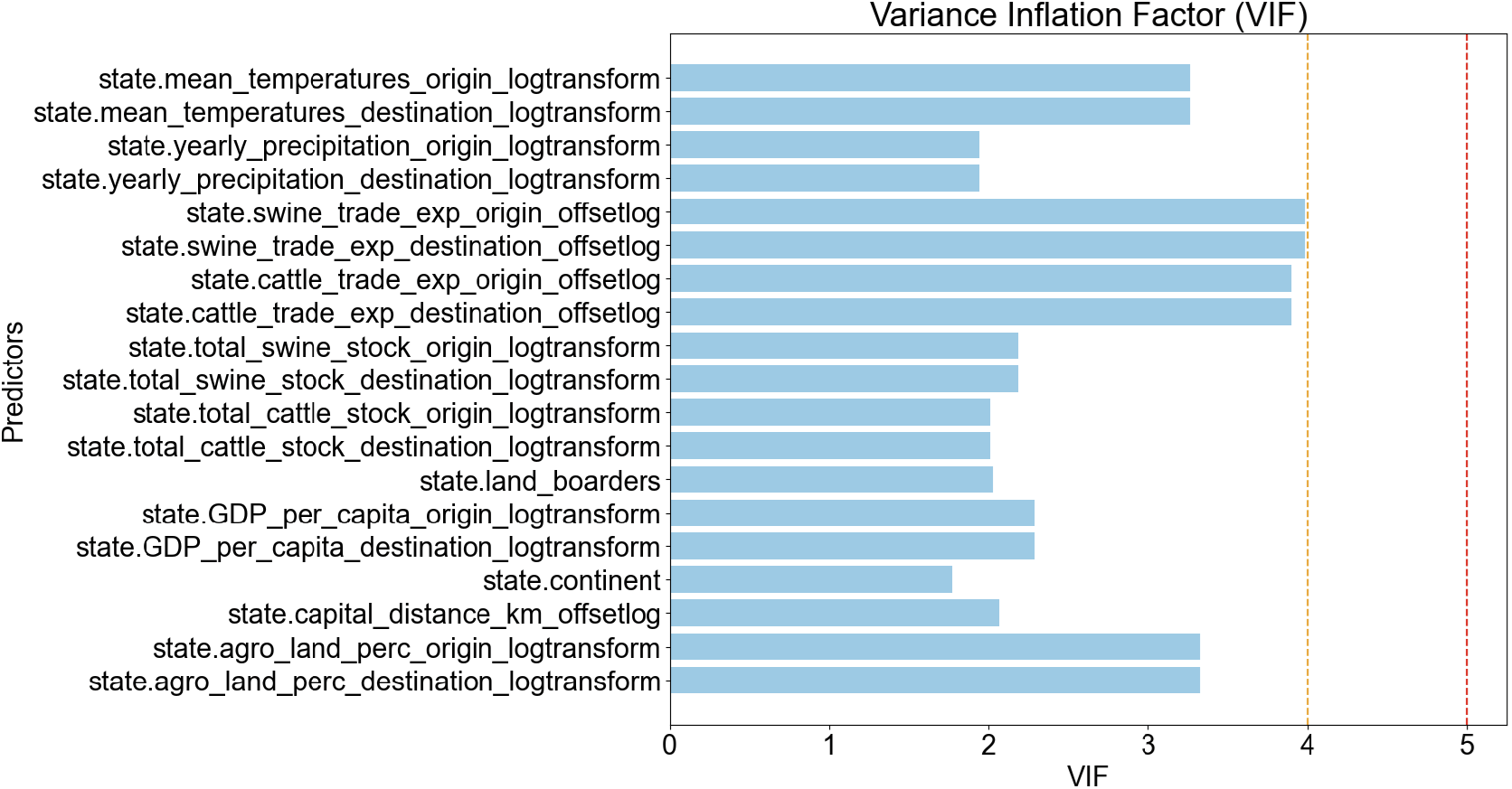
Colinearity of predictor variables (y-axis) after variable elimination. The common recommendations for upper variance inflation factor (VIF) limits of 4 and 5 are marked in orange and red, respectively (x-axis).

**Fig. S4:**
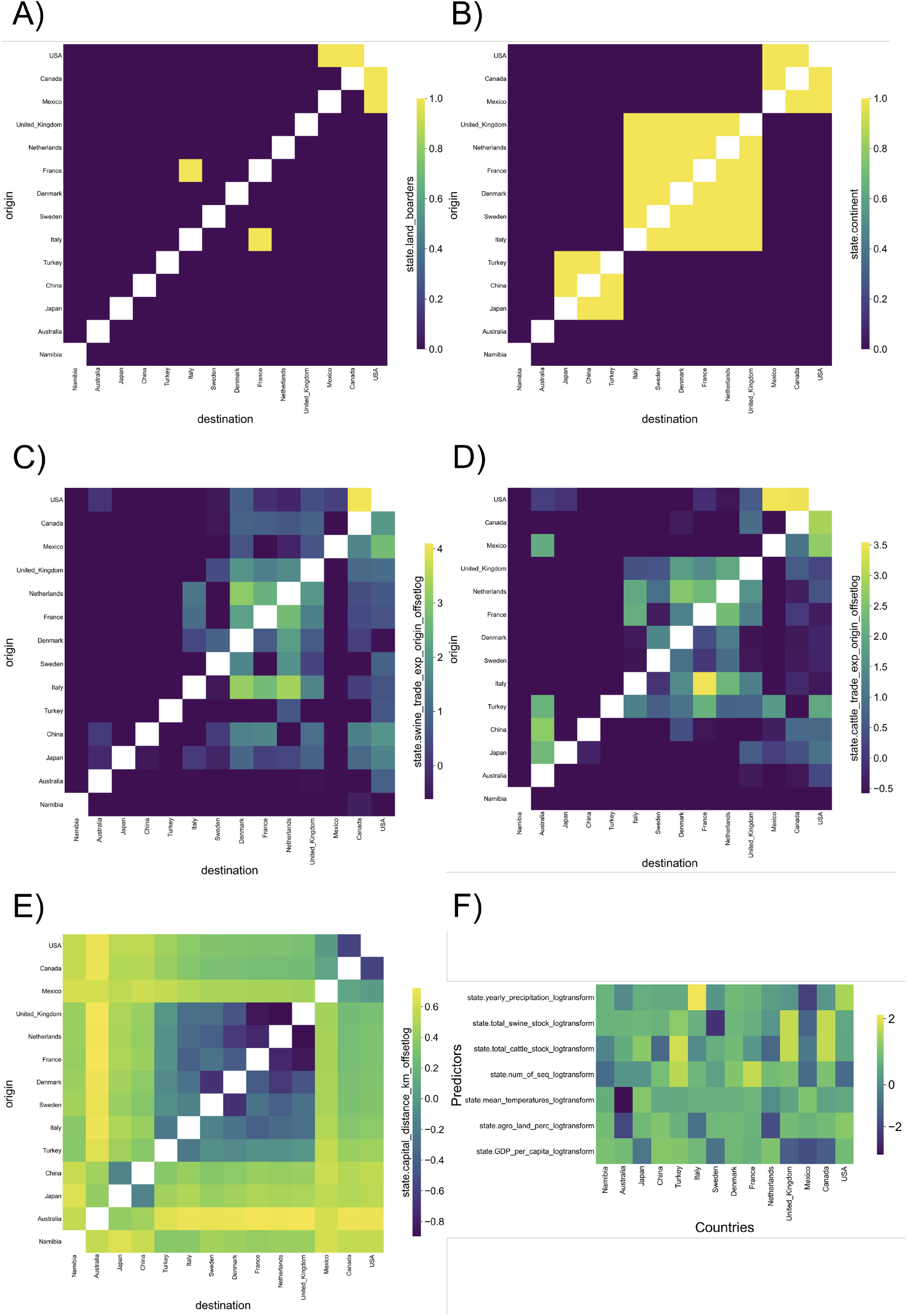
Heatmaps showing predictor matrices. A) Land borders dummy variable. B) Shared continent dummy variable. C) Swine trade standardized and log-transformed. D) Cattle trade standardized and log-transformed. E) distance between country capitals standardized and log-transformed. F) All other variables with single values per country (used to produce origin and destination predictors).

**Fig. S5:**
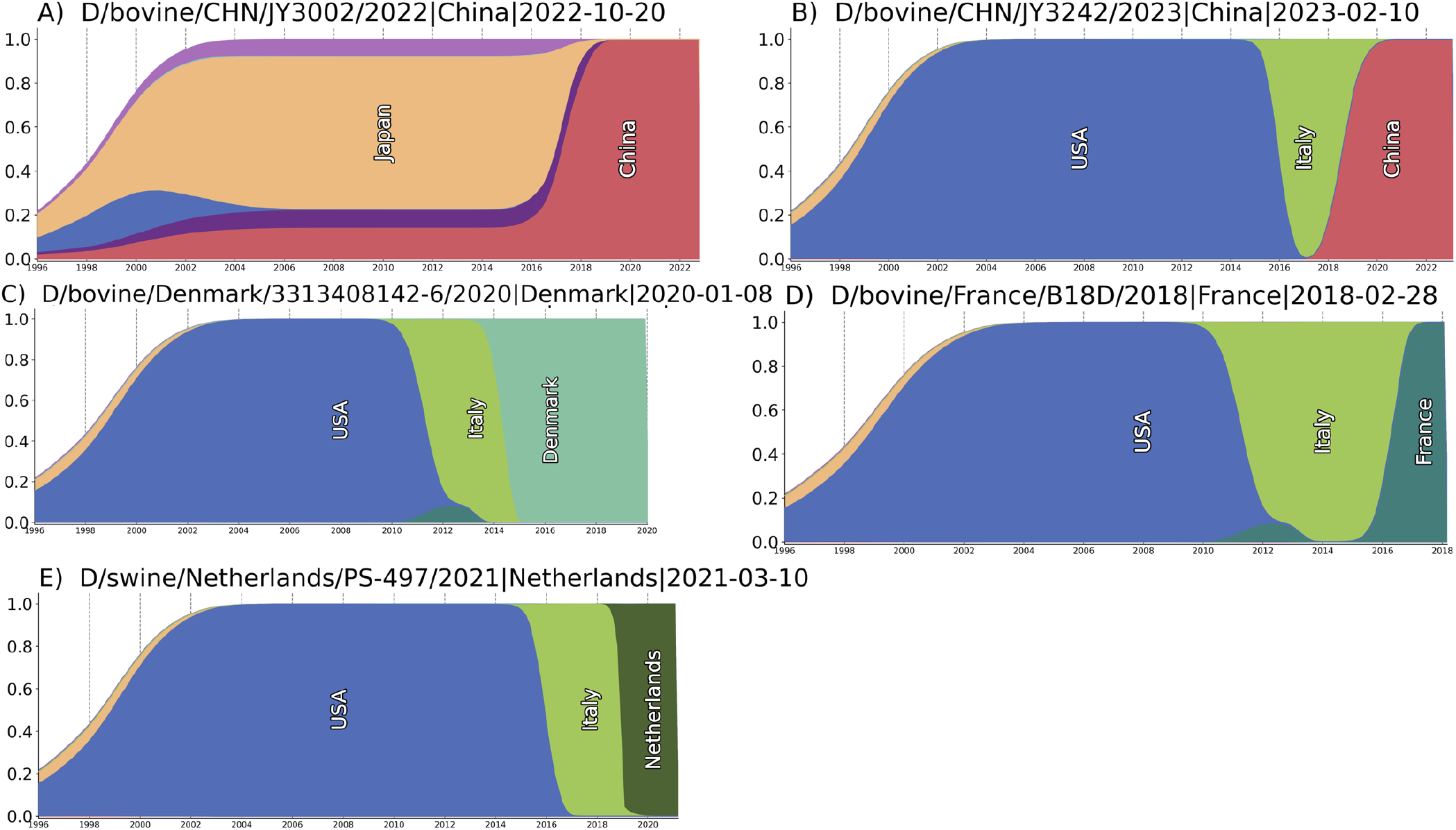
Individual state ancestry plots showing posterior probabilities of inferred ancestral states when tracing the path from the root of the tree to the following sequences: A) D/bovine/CHN/JY3002/2022 B) D/bovine/CHN/JY3242/2023 C) D/bovine/Denmark/3313408142-6/2020 D) D/bovine/France/B18D/2018 E) D/swine/Netherlands/PS-497/2021—Netherlands—2021-03-10.

**Fig. S6:**
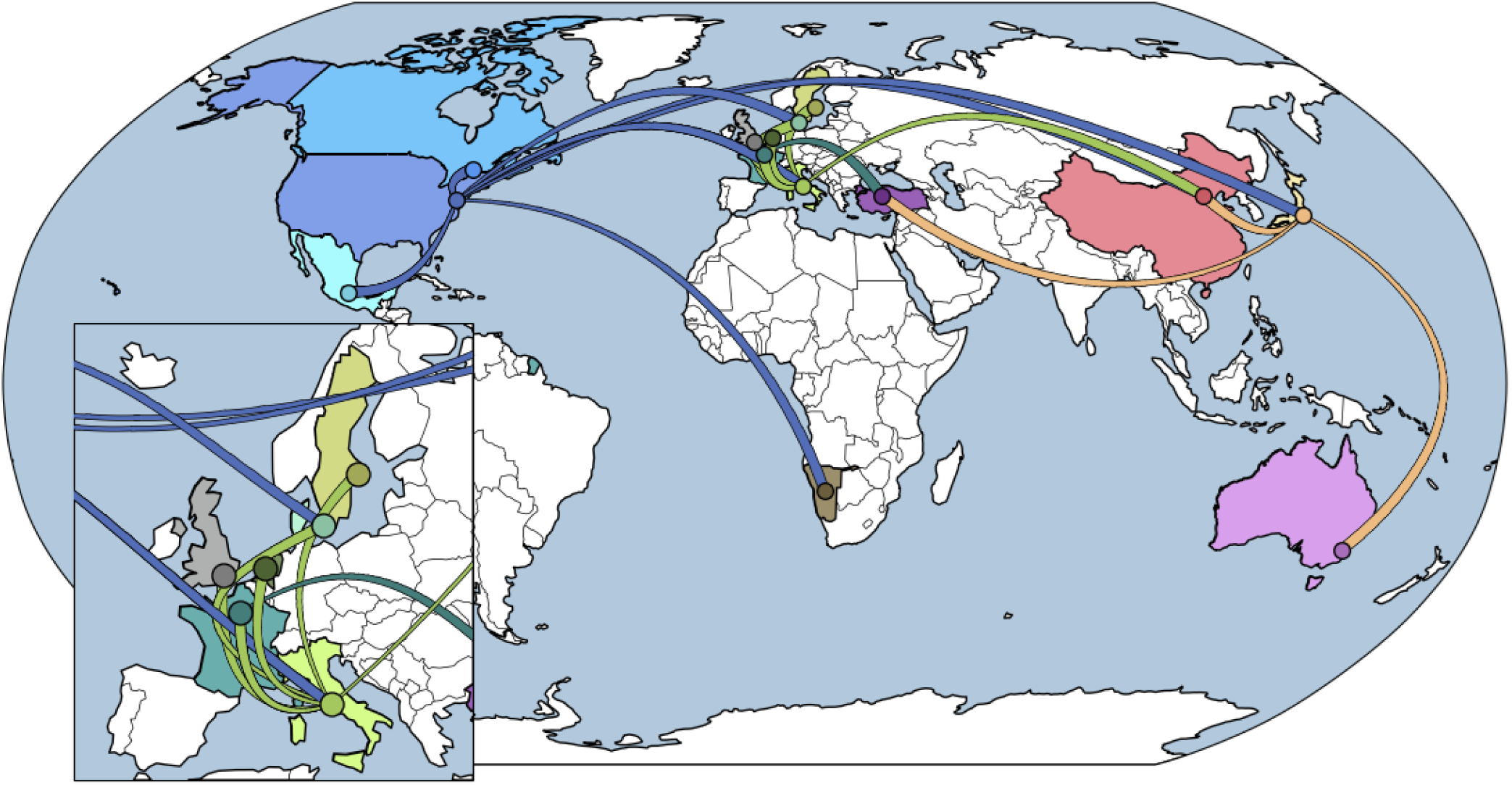
A global map showing inferred innfluenza D virus migrations. Countries with influenza D virus HEF sequence data that we used are coloured, with curved lines indicating inferred migration events (lines are thin at the origin location and widen towards the destination country).

